# Selection and validation of internal control genes for quantitative real-time RT–_q_PCR normalization of *Phlebopus portentosus* gene expression under different conditions

**DOI:** 10.1101/2022.10.19.512948

**Authors:** Chen-Menghui Hu, Jia-Ning Wan, Ting Guo, Guang-Yan Ji, Shun-Zhen Luo, Kai-Ping Ji, Yang Cao, Qi Tan, Da-Peng Bao, Rui-Heng Yang

## Abstract

*Phlebopus portentosus* (Berk. and Broome) Boedijin is an attractive edible mushroom and considered as the unique bolete to have achieved artificial cultivation *in vitro*. Gene expression analysis has become widely used in researches of edible fungi and is important to uncover the functions of genes involved in complex biological processes. Selecting appropriate reference genes is crucial to ensuring reliable results of gene expression analysis by RT–qPCR. In our study, a total of 12 candidate control genes were selected from 25 traditional housekeeping genes based on the expression stability in 9 transcriptomes of 3 developmental stages. These genes were further evaluated using *geNorm, NormFinder* and *RefFinder* under different conditions and developmental stages. The results revealed that MSF1-domain-containing protein (*MSF1*), synaptobrevin (*SYB*), mitogen-activated protein kinase genes (*MAPK*), TATA binding protein 1 (*TBP1*) and SPRY domain protein (SPRY) were the most stable reference genes in all sample treatments, while elongation factor 1-alpha (*EF1*), *Actin* and ubiquitin-conjugating enzyme (*UBCE*) were the most unstably expressed. The gene *SYB* was selected based on the transcriptome results and was identified as a novel reference gene in *P. portentosus*. This is the first detailed study for identification reference genes in this fungus and may provide new insights into selecting genes and quantifying gene expression.

## Introduction

*Phlebopus portentosus* (Berk. and Broome) Boedijin is an important ectomycorrhizal edible fungi and widely consumed in some regions of China, Thailand and some other tropical countries [1,2]. This fungus rich in nutrients [3,4] and has been successfully artificially cultivated in greenhouses on designed substrates *in vitro* in China and Thailand. It is the unique species with ectomycorrhizal niches in the order Boletales to achieve industrialized cultivation [1,5,6] and reaches production of 6 tons per day [1]. The success of artificial cultivation of this fungus plays important roles in domestications of wild fungi, especially for some ectomycorrhizal fungi. Gene expression analyses of fruiting-body formation, development and responses to environmental stresses are particularly important to understand the genetic backgroups in this fungus [7], answering the scientific questions on the functions of genes involved in complex biological and metabolic processes and the mechanisms of domestication [8–11]. Transcriptomes and reverse transcription quantitative real-time PCR (RT–qPCR) are the most commonly used techniques to determine the gene expression patterns of fruiting body development and lignocellulases synthesis in mushrooms [7]. However, up to now there are no related researches to explore the key steps affecting the results of RT–PCR in *P. portentosus*.

RT–qPCR is a highly sensitive tool used for the rapid and accurate characterization of gene expression in molecular experiments [9,12,13]. However, the accuracy and reliability of gene expression carried out using RT–qPCR is strongly influenced by lots of factors, such as the quality of raw samples, inhibitors included in samples, primer specificity, RNA quality and amount, efficiency of reverse transcription, internal reference genes and so on [11,14,15]. Of all the variations, the use of an optimal reference gene for RT–qPCR analysis is crucial to ensure accurate normalization, and unsuitable reference genes might cause unreliable and misleading results [16–18]. An ideal reference gene is characterized by a constant expression level across different conditions, including different environments and developments [13,19,20]. Traditionally, several traditional and housekeeping genes involved in some basic cellular and molecular processes have been widely used as reference genes, including 18S ribosomal RNA (*18S*), 28S ribosomal RNA (*28S*), *β*-actin (*ACTB*), cyclophilin (*CYP*), tubulin (*TUBα* and *TUBβ1*), glyceraldehyde-3-phosphate dehydrogenase (*GAPDH*), and ubiquitin (*UBQ*) [14,16,17,21–23]. Unfortunately, based on the results from animal and plant studies, no universal reference genes are suitable for all species in various tissues, multiple developmental stages or under different experimental treatments [19,20]. A similar situation also exists in mushroom studies, e.g., SPRY domain protein (*SPRY*), alpha-tubulin (*TUBα*), cyclophilin (*CYP*), Lasparaginase (*L-asp*), and MSF1-domain-containing protein (*MSF1*) were regarded as the most stably expressed genes under different conditions in *Volvariella volvacea* [16,21], while *UBQ* was the most stably expressed in *Auricularia cornea* [23]. Therefore, it is essential to validate suitable reference genes under different conditions to obtain biologically accurate expression data in special species.

There are a large number of published articles on selecting reference genes in different species, which reveal that the importance of control genes has been attached to RT-qPCR normalization. In addition to the two species above (*Volvariella volvacea* and *Auricularia cornea*), some other studies have also been performed to identify and evaluate reference genes in mushrooms, including *Agaricus bisporus* (J.E. Lange) Imbach [24], *Ganoderma lucidum* (Curtis) P. Karst. [25], *Pleurotus ostreatus* (Jacq.) P. Kumm [26], *Ophiocordyceps sinensis* (Berk.) G.H. Sung, J.M. Sung, Hywel-Jones & Spatafora [27], *Wolfiporia cocos* (F.A. Wolf) Ryvarden & Gilb [28,29], *Auricularia heimuer* F. Wu, B.K. Cui & Y.C. Dai [30], *Lepista sordida* (Schumach.) Singer [31], *Inonotus oblisquus* [32], *Flammulina filiformis* (Z.W. Ge, X.B. Liu & Zhu L. Yang) P.M. Wang, Y.C. Dai, E. Horak & Zhu L. Yang [32–34], *Macrocybe gigantea* (Massee) Pegler & Lodge [35], *Armillaria mellea* (Vahl) P. Kumm [36], *Pleurotus eryngii* (DC.) Quél and [37], *Tuber melanosporum* Vittad [38] and *Morchella importuna* M. Kuo, O’Donnell & T.J. Volk [39]. All the results also revealed that different reference genes are suitable for different species and different conditions. However, studies of the stability and identification of reference genes have not been carried out in *P. portentosus*.

It is difficult to detect the optimal reference genes in a species without genomic and transcriptomic information [39–41]. Fortunately, there are some related datasets deposited in the NCBI database [7,42], which will be convenient for screening suitable reference genes and primer designs for *P. portentosus*. In this study, some candidate reference genes were pre-evaluated and selected based on genomic and transcriptomic data. The expression patterns of filtered candidate reference genes under different treatments were further validated. Our results will provide new insights into reference gene selection for high accuracy of RT-qPCR normalization in *P. portentosus* gene expression analysis.

## Materials and Methods

### Sampling and culture conditions

The *P. portentosus* dikaryon strain 17026 used in this study was the same as used in our previous work [7], which provided by Hongzhen Agricultural Science and Technology Co. Ltd. and maintained at the Shanghai Academy of Agricultural Science. This strain was incubated in potato dextrose agar (PDA) at 30 °C under dark as previously described [7,42]. Mycelial samples were prepared in PDA medium for 7 days under selected conditions with shaking (150 rpm). Addition of NaCl (1%, 30 °C), CuSO_4_ (1%, 30 °C), H_2_O_2_ (100 μM, 30 °C), HCl (pH 4.0, 30 °C), NaOH (pH 9.0, 30 °C) created conditions of salt, heavy metal, oxidation, acid and alkali stresses. The optimal temperature was 30 °C for this fungus, incubated at 25 °C and 35 °C to create cold and heat conditions respectively. The artificial cultivation of *P. portentosus* was carried out in line with the methods published in our previous study [7]. The entire fruiting body was harvested, and cap and stipe were chopped into small pieces (2 mm) . All samples were immediately frozen in liquid nitrogen and stored at −80 °C before RNA extraction. Three independent biological replicates were tested for each sample.

### RNA isolation and cDNA synthesis

A total of 100 mg samples of mycelia were grinded into power using liquid nitrogen and then treated with TRIzol (Invitrogen) and DNase I (Ambion, USA) to isolated RNA and remove contaminating DNA. The procedure of RNA purification followed the manufacturer’s protocol. RNA concentration and purity were assessed using a NanoDrop 2000 Spectrophotometer (NanoDrop Technologies, Thermo Scientific, USA). Only RNA preparations with absorption ratio values of A260/280 and A260/A230 were 1.8-2.0 and >2.0 were used for subsequent analysis. cDNA was synthesized using the PrimeScript™ RT reagent Kit with gDNA Eraser (Perfect Real Time) (TaKaRa Bio Inc., Dalian, China) from 1 *μ*g total RNA in a final volume of 20 *μ*L according to the provided protocol . The cDNA obtained were diluted 10 fold using nuclease-free water for further RT–qPCR amplifications.

### Selection and validation of candidate reference genes and primer design

Based on previous literatures, a total of 25 housekeeping genes [14,16,17,21–24,26–30,33–35,37–39,43], e.g., 60S ribosomal protein (*60S*), Ras protein (*RAS*), actin 2 (*ACTIN*), adenine phosphoribosyl transferase (*APT*), cyclophilin (*CYP*), 18S ribosomal RNA (*18S* rRNA), RNA polymerase subunit 2 (*POL II*), RNA polymerase subunit 3 (*POL III*), elongation factor 1-alpha (*EF1*), elongation factor 2-alpha (*EF2*), eukaryotic initiation factor 4A (*EIF*), glyceraldehyde-3-phosphate dehydrogenase (*GAPDH*), GTP-binding nuclear protein (*RAN*), ribosomal protein S (RPS), s-adenosyl methionine decarboxylase (SAMDC), TATA binding protein 1 (*TBP1*), *alpha*-tubulin (*TUBα*), *beta*-Tubulin (*TUBβ*), ubiquitin-conjugating enzyme (*UBCE*), ubiquitin (*UBQ*), polyubiquitin (POL), MSF1-domain-containing protein (*MSF1*), SPRY domain protein (*SPRY*), L-asparaginase (*ASP*), and mitogen-activated protein kinase genes (*MAPK*), were selected. All candidate genes were pre-evaluated using transcriptome datasets from our previous study on developmental stages [21]. Gene expression was normalized using the fragments per kilobase of transcript million mapped reads (FPKM)>1 (FPKM) method [21,43–47]. The average expression (mean) and the standard deviation (SD) were calculated per gene over the entire set of transcriptome data. Genes with low CV values (SD/mean) were considered to own high expression stability than genes with high CV values. The cutoff CV value for stable gene expression was often set as <0.3 in line with the studies published previously [46,48]. The primer pairs were designed using Blast-Primer based on the following criteria: amplified products ranging from 100-150bp, Tm (melting temperature) ranging from 55 to 60 °C, GC content ranging from 15% to 55%, and primer length ranging from 20 to 25.

### RT–qPCR and data analysis

RT-qPCR was carried out on an Applied Biosystems 7500 Real-Time PCR system, and samples were run in triplicate. Each PCR reaction mixtures were prepared in a final volume of 20 μL contained 2 *μ*l of prepared cDNA template (10 fold diluted template), 0.4 *μ*l of each primer (10 nM, reverse and forward primers), 6.8 *μ*l of ddH_2_O and 10 *μ*l of Power SYBR Green qPCR Master Mix (TaKaRa). Three technical replicates, including no-template controls, were used for each sample. The running conditions and parameter used were as follows: initial denaturation at 95 °C for 5 min, followed by 40 cycles of 95 °C for 15 s and 60 °C for 1 min. A temperature ramp step was used for dissociation analysis to confirm the single product with heating from 60°C to 95°C.

To analyze the expression stability of candidate reference genes, three different programs, *geNorm* [49], *NormFinder* [50], and *BestKeeper* [51], were used based on the experimental design and manufacturers’ instructions. The detailed analysis methodology followed the previous methods [16].

## Results

### pre-evaluation of candidate reference genes based on transcriptome data

In order to identify stably expressed genes in *P. portentosus*, a total of 9 transcriptomes collected from 3 developmental stages, including mycelium, primordium and fruiting body, were used. Based on the CV value of gene expression (RPKM) and the cutoff for stably expressed genes (CV < 0.3) between different stages [46], 952 genes were stably expressed in all the samples. Based on the annotation of genes, synaptobrevin (*SYB*) was the most stable in the three stages with CV=0.034. Among the 25 candidate genes above, 11 genes were detected to be stably expressed in development stages with CV values ranging from 0.15 to 0.30 (Fig. 1), including *MSF1, RAN, ACTIN, UBCE, MAPK, SPRY, EF1, EF2, CYP, TBP1* and *RP*. Some studies revealed that the genes with CV < 0.35 were also accepted as reference candidates [27]. Finally, 12 candidate genes and *SYB* were used for RT–PCR validation.

**Fig. 1.**
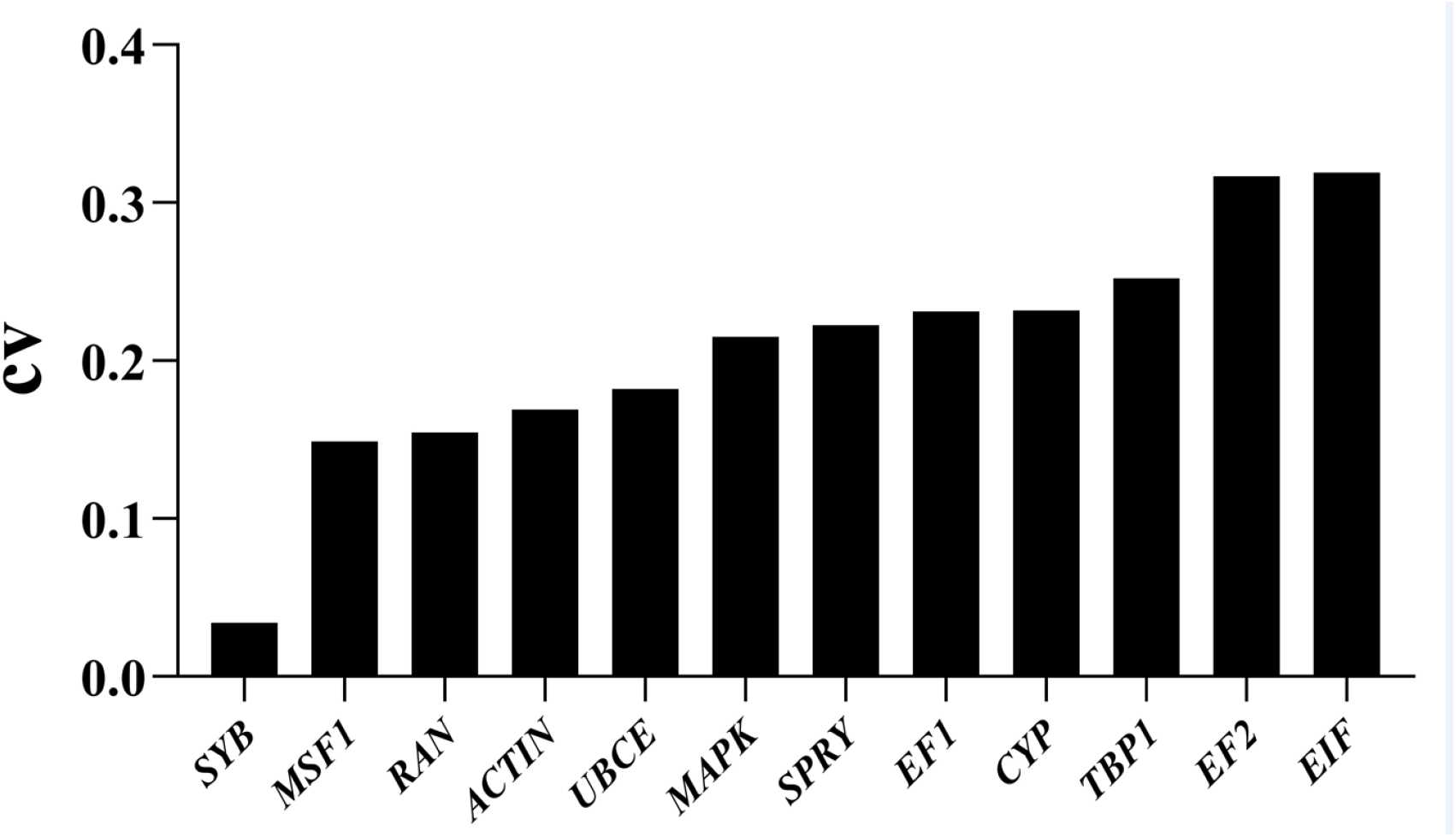
CV values of the RPKMs of 12 candidate reference genes from the transcriptome datasets

### Specificity of amplification and PCR efficiency

Information on the primer pairs is listed in Table 1. To confirm the specificity of the primers, 2% agarose gel electrophoresis was used. PCR products amplified using each primer pair on cDNA samples with the expected size, a single band and without primer dimers were considered to be high specificity (Fig. S1).

**Table 1.**
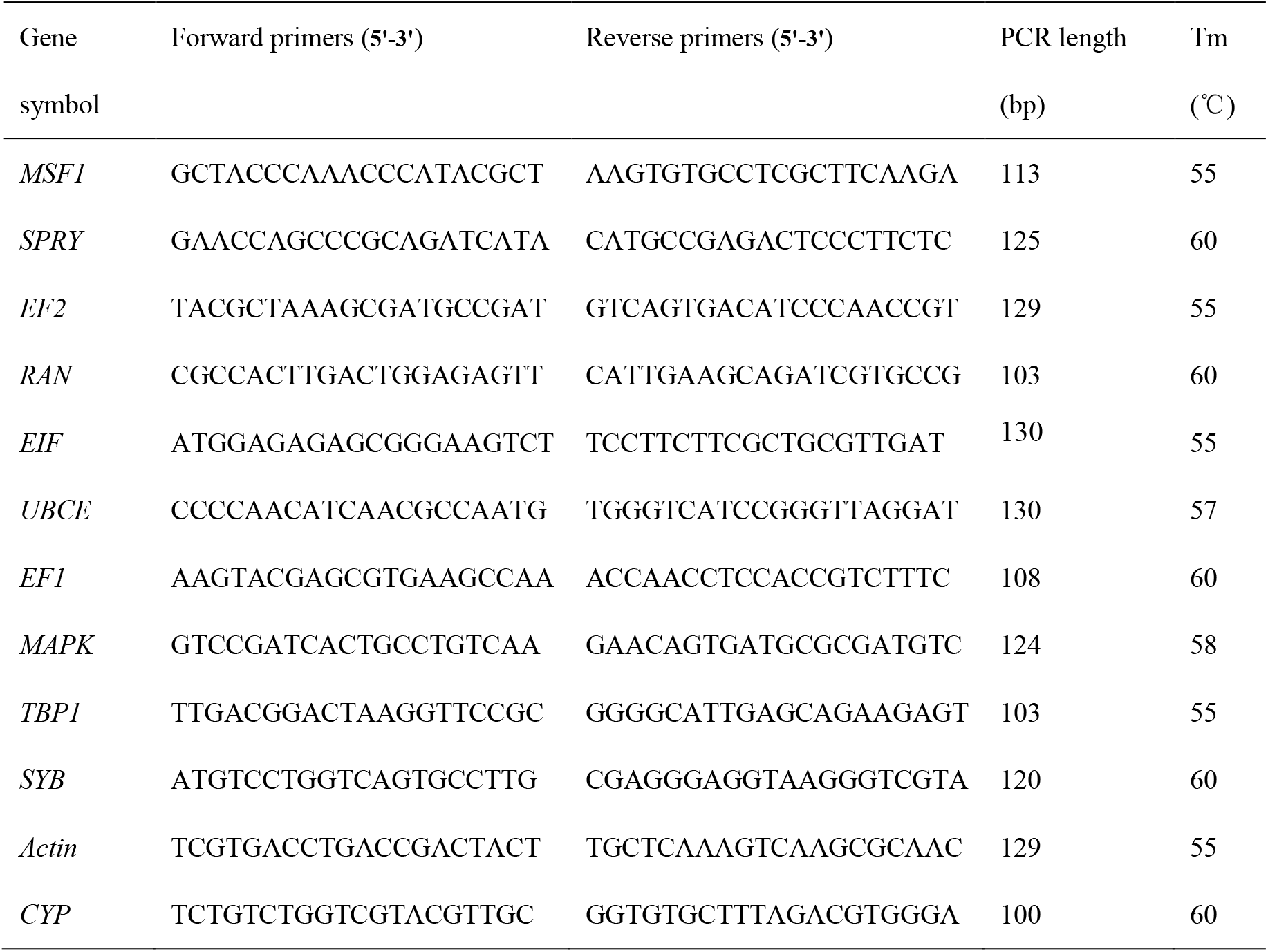
The information of selected candidate reference genes, primers, and amplicon length.

### Reference gene candidate expression profiles

To evaluate the expression levels of the 12 selected reference genes from the transcriptomic analysis, threshold cycle (Ct) values were calculated from the total samples. The average Ct values for the 12 genes ranged from 13.45 to 30.31, and most of Ct values were between 21 and 28. *UBCE* was the most abundantly expressed gene, and *SYB* was the lowest expressed gene.

### *geNorm* analysis

The analysis of *geNorm* was based on the ‘pairwise comparison strategy’, the gene with the lowest M value the most stable expression and would be selected as potential reference genes, while the gene with the highest M value had greater variation in expression. The result shown in Fig. 2 revealed that different reference genes had different stabilities under different conditions. The top 2 stable reference genes for RT– qPCR normalization were *TBP1* and *RAN* for the fruiting body stage (M<0.5), *ACT* and *EIF* for heat stress and heavy metal stress (M<0.5), *TBP1* and *MAPK* for cold stress (M<0.5), EF2 and TBP1 for oxidative stress (M<0.5), *ACT* and *EF2* for salt stress (M<0.5), *SYB* and *SPRY* for acid stress (M<0.5), *MAPK* and *RAN* for alkali stress (M<0.5). Except for *UBCE*, all the genes had the highest M values <1.0 across all samples, and *SYB* and *SMF1* were the most stable genes (Fig. 2). Therefore, these two reference genes were regarded as the best reference genes for the widely used under test conditions based on our data.

**Fig. 2.**
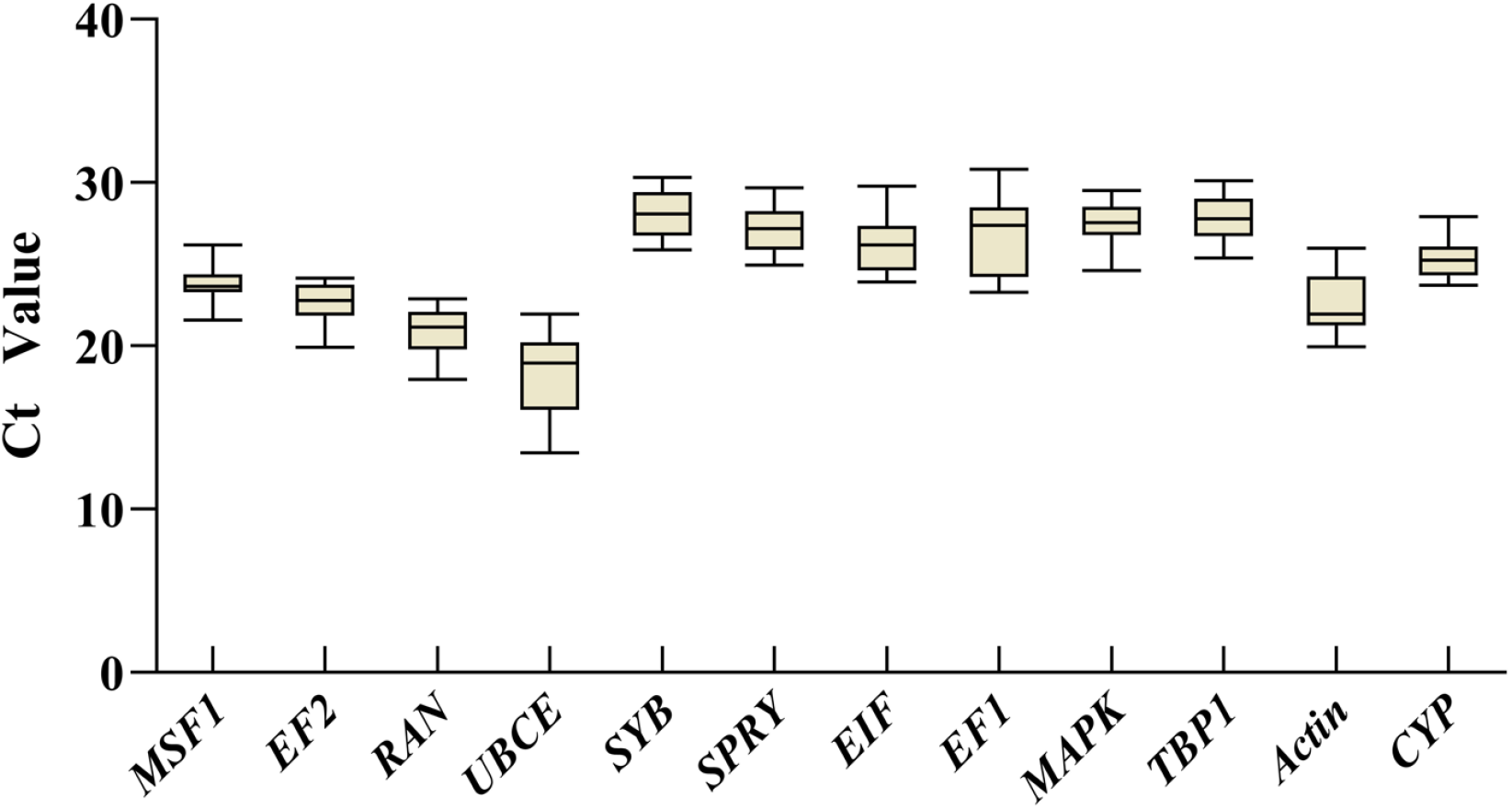
Distribution overview of threshold cycle (Ct) values for candidate reference genes across tested samples in *P. portentosus*. The lines crossing the boxes represent the medians. The box plots represent the 25th and 75th percentiles. The whiskers represent the minimum and maximum values.

It was acknowledged that combination of two or more reference genes in RT–qPCR analysis might generate high accuracy results than single gene. The pairwise variations Vn/Vn + 1 between two normalization factors (NF values NFn, where n = number of genes included) calculated using *geNorm* software could estimate the optimal number of genes required for accurate normalization. the results showed that all the V-values were < 0.15 (Fig. 4). However, the average *geNorm* M-value of genes (Fi. 1) in all the treatments was not lower than 0.2, which suggested that only two reference genes were adequate for normalizing gene expression data.

**Fig. 3.**
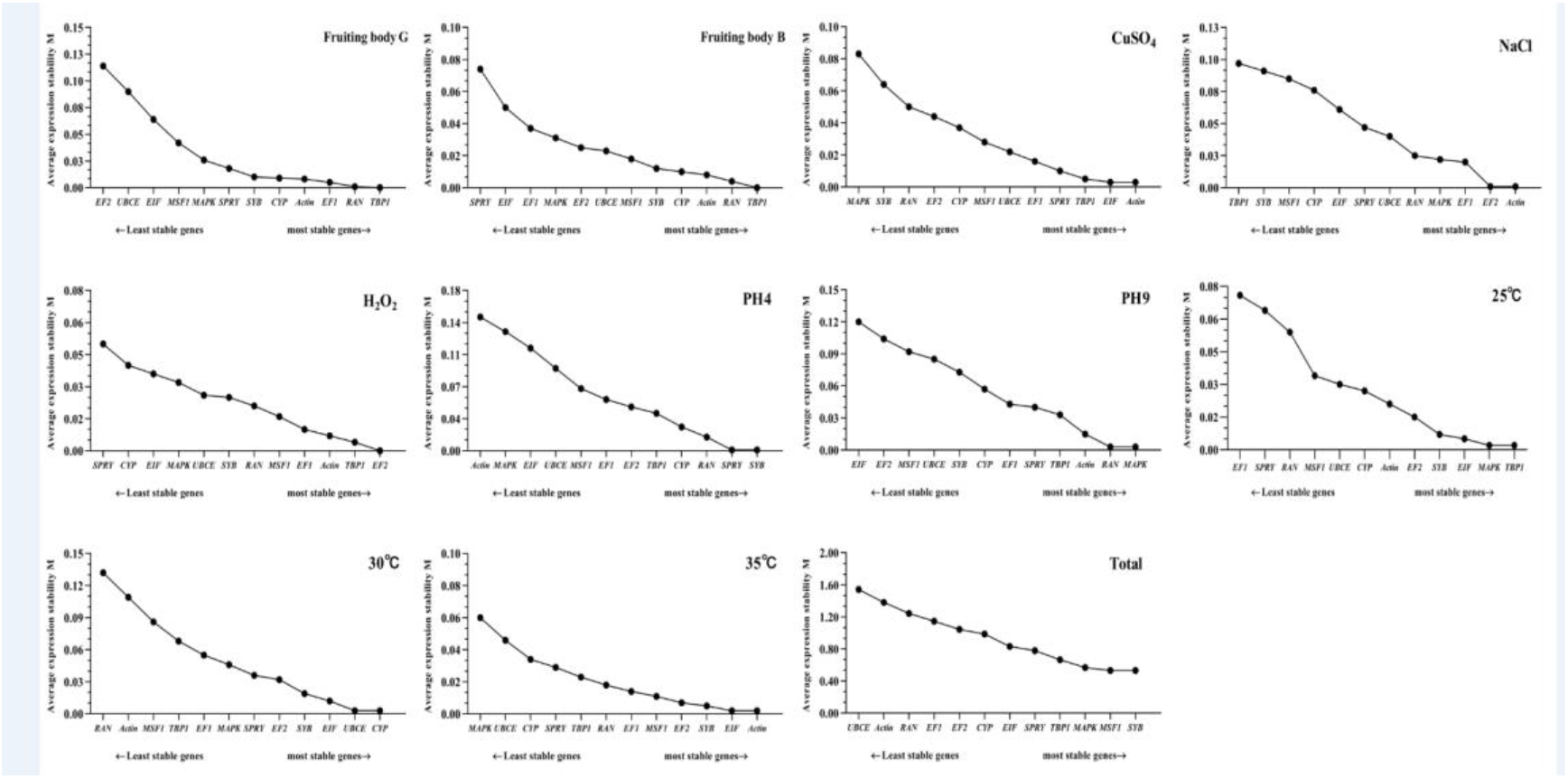
Stability of the 12 candidate reference genes calculated by geNorm under different conditions. The average expression stability values (M) of the reference genes were measured. The least stable gene with the highest M value is located on the left, while the most stable gene is located on the right.

**Fig. 4.**
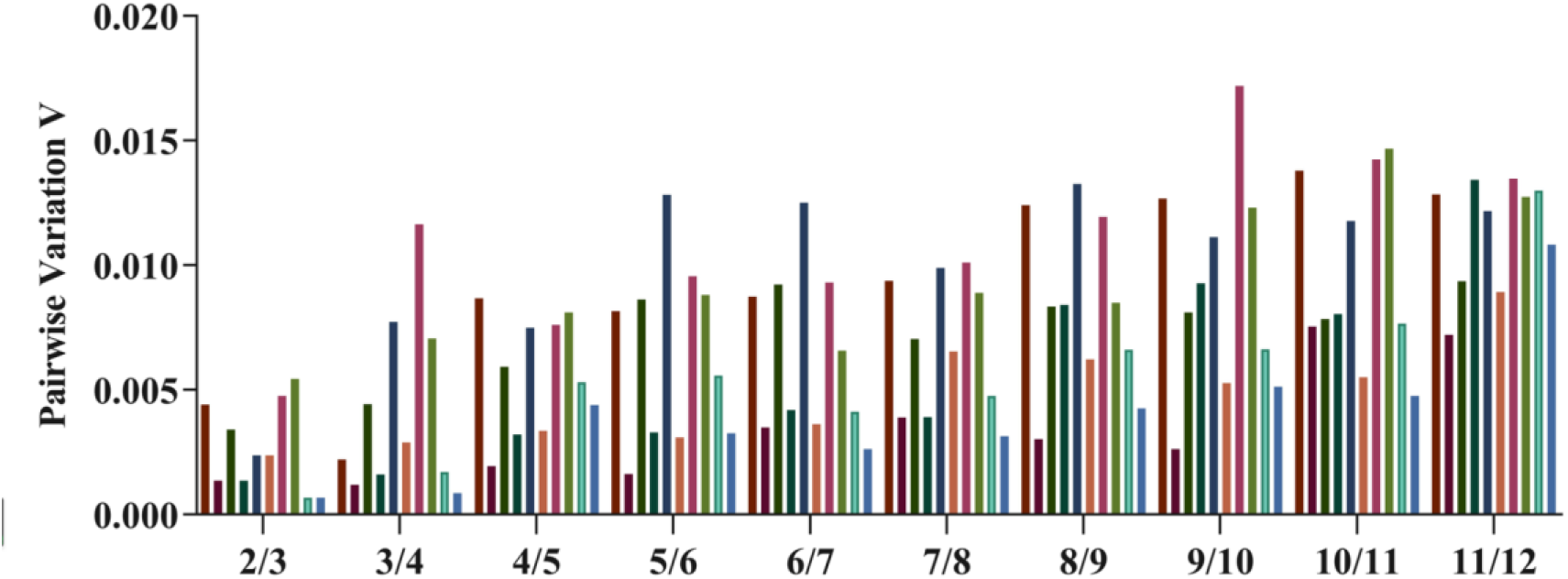
Determination of the optimal number of reference genes for normalization. Pairwise variations (Vn/n+1) calculated using geNorm. V is the variation value, where >0.15 indicates that an additional reference gene does not improve normalization.

### *NormFinder* Analysis

The program *NormFinder* takes into account the two most stable reference genes in intra- and intergroup expression, which was used to evaluate the optimal reference genes for RT–qPCR normalization. A lower stability value indicated a higher stability. The gene expression stabilities evaluated using *NormFinder* was listed in Table 2. The ranking order from the top to the bottom revealed that the stabilities decreased. More than 10 genes were candidate genes with stability values (SV) <0.15 during development, and *Actin* and *CYP* were the most stable (SV=0.00). *Actin, EIF*, and *TBP1* were the most stable under heavy metal stress (SV=0.002), *EF1* and *Actin* were the most stable under salty stress (SV=0.003), *EF2* and *MAPK* were most stable under acid stress, *TBP1* and *RAN* were the most stable under alkali stress, *CYP* and *UBCE* were the most stable under cold stress, and *Actin* and *EIF* were the most stable under heat stress. Across all the samples, MSF1 was the most stable, followed by *SYB, MAPK* and *EF2*.

**Table 2.** Expression stability of the 12 selected reference genes evaluated by *NormFinder*

| Rank | G | B | CuSO<sub>4</sub> | NaCl | H<sub>2</sub>O<sub>2</sub> | PH4 | PH9 | 25°C | 30°C | 35°C | Total |
| --- | --- | --- | --- | --- | --- | --- | --- | --- | --- | --- | --- |
| 1 | *Actin* | *Actin* | *Actin* | *SPRY* | *EF1* | *EF2* | *MAPK* | *CYP* | *SPRY* | *Actin* | *MSF1* |
|  | 0.000 | 0.001 | 0.002 | 0.003 | 0.003 | 0.003 | 0.001 | 0.002 | 0.000 | 0.001 | 0.158 |
| 2 | *CYP* | *CYP* | *EIF* | *UBCE* | *Actin* | *TBP1* | *RAN* | *UBCE* | *EF2* | *EIF* | *SYB* |
|  | 0.000 | 0.001 | 0.002 | 0.004 | 0.003 | 0.003 | 0.001 | 0.002 | 0.000 | 0.001 | 0.386 |
| 3 | *TBP1* | *MAPK* | *TBP1* | *EIF* | *MSF1* | *EF1* | *Actin* | *Actin* | *SYB* | *SYB* | *MAPK* |
|  | 0.000 | 0.008 | 0.002 | 0.039 | 0.006 | 0.031 | 0.01 | 0.005 | 0.010 | 0.002 | 0.409 |
| 4 | *RAN* | *TBP1* | *UBCE* | *RAN* | *EF2* | *SPRY* | *TBP1* | *EF2* | *UBCE* | *EF2* | *EF2* |
|  | 0.001 | 0.016 | 0.006 | 0.061 | 0.014 | 0.032 | 0.042 | 0.006 | 0.017 | 0.002 | 0.636 |
| 5 | *EF1* | *RAN* | *MSF1* | *MAPK* | *RAN* | *SYB* | *CYP* | *MSF1* | *CYP* | *MSF1* | *TBP1* |
|  | 0.003 | 0.025 | 0.017 | 0.074 | 0.021 | 0.034 | 0.054 | 0.017 | 0.023 | 0.011 | 0.751 |
| 6 | *SYB* | *EF1* | *SPRY* | *EF1* | *TBP1* | *RAN* | *SPRY* | *SYB* | *EIF* | *EF1* | *SPRY* |
|  | 0.004 | 0.03 | 0.021 | 0.08 | 0.022 | 0.059 | 0.059 | 0.038 | 0.048 | 0.022 | 0.983 |
| 7 | *MAPK* | *SYB* | *EF1* | *CYP* | *SYB* | *CYP* | *EF1* | *TBP1* | *MAPK* | *TBP1* | *CYP* |
|  | 0.017 | 0.033 | 0.038 | 0.094 | 0.031 | 0.092 | 0.068 | 0.045 | 0.101 | 0.025 | 1.113 |
| 8 | *SPRY* | *MSF1* | *CYP* | *MSF1* | *UBCE* | *MSF1* | *SYB* | *MAPK* | *MSF1* | *RAN* | *EIF* |
|  | 0.049 | 0.058 | 0.057 | 0.101 | 0.032 | 0.143 | 0.126 | 0.048 | 0.121 | 0.032 | 1.137 |
| 9 | *MSF1* | *UBCE* | *EF2* | *SYB* | *CYP* | *EIF* | *EF2* | *EIF* | *EF1* | *SPRY* | *RAN* |
|  | 0.125 | 0.069 | 0.068 | 0.108 | 0.05 | 0.168 | 0.143 | 0.054 | 0.126 | 0.046 | 1.234 |
| 10 | *EIF* | *EF2* | *RAN* | *Actin* | *MAPK* | *MAPK* | *UBCE* | *RAN* | *TBP1* | *CYP* | *EF1* |
|  | 0.199 | 0.069 | 0.072 | 0.112 | 0.062 | 0.192 | 0.146 | 0.079 | 0.169 | 0.053 | 1.766 |
| 11 | *UBCE* | *EIF* | *SYB* | *EF2* | *EIF* | *UBCE* | *MSF1* | *SPRY* | *Actin* | *UBCE* | *Actin* |
|  | 0.200 | 0.093 | 0.115 | 0.113 | 0.065 | 0.23 | 0.153 | 0.101 | 0.196 | 0.12 | 1.802 |
| 12 | *EF2* | *SPRY* | *MAPK* | *TBP1* | *SPRY* | *Actin* | *EIF* | *EF1* | *RAN* | *MAPK* | *UBCE* |
|  | 0.242 | 0.209 | 0.188 | 0.125 | 0.107 | 0.233 | 0.208 | 0.109 | 0.256 | 0.137 | 2.242 |

### *BestKeeper* analysis

*BestKeeper* is an Excel-based tool based on pairwise correlation analysis of selected reference genes through calculating the standard deviation (SD) and coefficient of variance (CV), which could be used to help in selection of suitable reference genes. The lowest CV and SD was considered as the most stably expressed evaluation criteria; gradual increases suggested the stability decreased. It is essential to note that genes with an SD > 1 should be considered unacceptable. The rankings of *BestKeeper* analysis revealed that *SYB* was the most suitable for samples in cold and heavy metal environments, *EIF* in a heat environment, *MSF1* in an oxidative environment, *UBCE* in a salt environment, *SPYR* in an acid environment, and *Actin* in an alkali environment (Fig. 5). *MSF1* and *CYP* were the most stable genes with the highest correlation coefficient. These results were in agreement with those selected by *geNorm* and *NormFinder*.

**Fig. 5.**
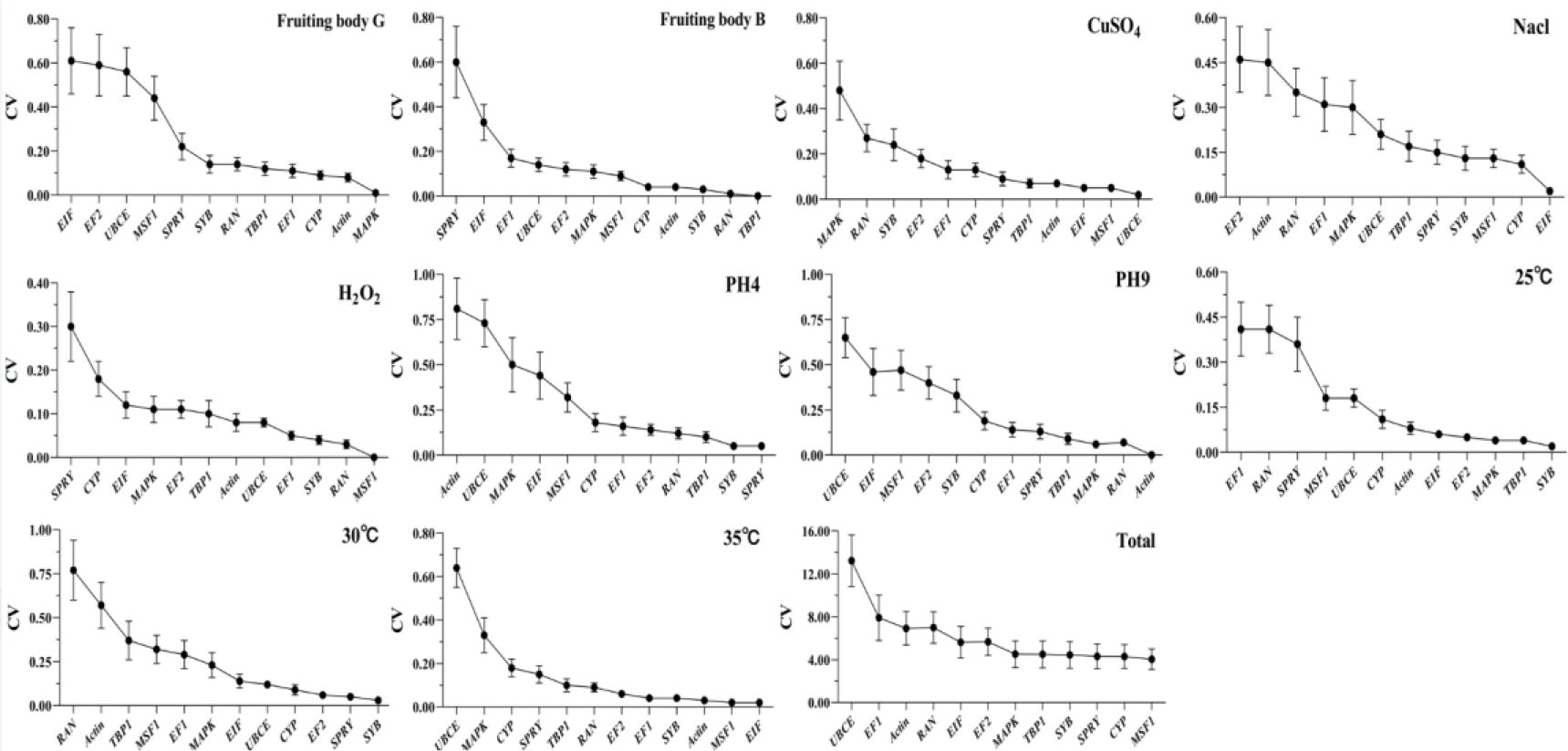
Stability of the 12 candidate reference genes evaluated by BestKeeper under different conditions. The lowest CV and SD revealed the most stable locating on the right of the plot, while the least stable gene locating on the left.

### Comprehensive stability analysis of reference genes

In order to get a consensus result from the three methods above, the geometric means of combinations of all the results and the corresponding rankings for each candidate gene were used (Table 3). The combination results revealed that *MSF1, SYB, MAPK, TBP1* and *SPRY* were the top 5 stable reference genes in all sample treatments. In contrast, *EF1, Actin* and *UBCE* were unstably expressed in most of tested samples. Based on geometric means of the three algorithms’ corresponding rankings, the results were more intuitive.

**Table 3.** Expression stability ranking of the 12 candidate reference genes.

| Method | 1 | 2 | 3 | 4 | 5 | 6 | 7 | 8 | 9 | 10 | 11 | 12 |
| --- | --- | --- | --- | --- | --- | --- | --- | --- | --- | --- | --- | --- |
| Genorm | *TBP1* | *RAN* | *EF1* | *Actin* | *CYP* | *SYB* | *SPRY* | *MAPK* | *MSF1* | *EIF* | *UBCE* | *EF2* |
| NormFinder | *Actin* | *CYP* | *TBP1* | *RAN* | *EF1* | *SYB* | *MAPK* | *SPRY* | *MSF1* | *EIF* | *UBCE* | *EF2* |
| Bestkeeper | *MAPK* | *Actin* | *CYP* | *EF1* | *TBP1* | *RAN* | *SYB* | *SPRY* | *MSF1* | *UBCE* | *EF2* | *EIF* |
| Comprehensive ranking | *TBP1* | *Actin* | *CYP* | *RAN* | *EF1* | *MAPK* | *SYB* | *SPRY* | *MSF1* | *EIF* | *UBCE* | *EF2* |
| Genorm | *TBP1* | *RAN* | *Actin* | *CYP* | *SYB* | *MSF1* | *UBCE* | *EF2* | *MAPK* | *EF1* | *EIF* | *SPRY* |
| NormFinder | *Actin* | *CYP* | *MAPK* | *TBP1* | *RAN* | *EF1* | *SYB* | *MSF1* | *UBCE* | *EF2* | *EIF* | *SPRY* |
| Bestkeeper | *TBP1* | *RAN* | *SYB* | *Actin* | *CYP* | *MSF1* | *MAPK* | *EF2* | *UBCE* | *EF1* | *EIF* | *SPRY* |
| Comprehensive ranking | *TBP1* | *Actin* | *RAN* | *CYP* | *SYB* | *MAPK* | *MSF1* | *UBCE* | *EF2* | *EF1* | *EIF* | *SPRY* |
| Genorm | *EIF* *Actin* |  | *TBP1* | *SPRY* | *EF1* | *UBCE* | *MSF1* | *CYP* | *EF2* | *RAN* | *SYB* | *MAPK* |
| NormFinder | *Actin* | *EIF* | *TBP1* | *UBCE* | *MSF1* | *SPRY* | *EF1* | *CYP* | *EF2* | *RAN* | *SYB* | *MAPK* |
| Bestkeeper | *UBCE* | *MSF1* | *EIF* | *Actin* | *TBP1* | *SPRY* | *CYP* | *EF1* | *EF2* | *SYB* | *RAN* | *MAPK* |
| Comprehensive ranking | *EIF* | *Actin* | *UBCE* | *TBP1* | *MSF1* | *SPRY* | *EF1* | *CYP* | *EF2* | *RAN* | *SYB* | *MAPK* |
| Genorm | *EF2* *Actin* |  | *EF1* | *MAPK* | *RAN* | *UBCE* | *SPRY* | *EIF* | *CYP* | *MSF1* | *SYB* | *TBP1* |
| NormFinder | *SPRY* | *UBCE* | *EIF* | *RAN* | *MAPK* | *EF1* | *CYP* | *MSF1* | *SYB* | *Actin* | *EF2* | *TBP1* |
| Bestkeeper | *EIF* | *CYP* | *MSF1* | *SYB* | *SPRY* | *TBP1* | *UBCE* | *MAPK* | *EF1* | *RAN* | *Actin* | *EF2* |
| Comprehensive ranking | *SPRY* | *EIF* | *UBCE* | *RAN* | *CYP* | *MAPK* | *Actin* | *EF1* | *EF2* | *MSF1* | *SYB* | *TBP1* |
| Genorm | *EF2* | *TBP1* | *Actin* | *EF1* | *MSF1* | *RAN* | *SYB* | *UBCE* | *MAPK* | *EIF* | *CYP* | *SPRY* |
| NormFinder | *EF1* | *Actin* | *MSF1* | *EF2* | *RAN* | *TBP1* | *SYB* | *UBCE* | *CYP* | *MAPK* | *EIF* | *SPRY* |
| Bestkeeper | *MSF1* | *RAN* | *SYB* | *EF1* | *UBCE* | *Actin* | *TBP1* | *EF2* | *MAPK* | *EIF* | *CYP* | *SPRY* |
| Comprehensive ranking | *EF1* | *MSF1* | *Actin* | *EF2* | *RAN* | *TBP1* | *SYB* | *UBCE* | *MAPK* | *CYP* | *EIF* | *SPRY* |
| Genorm | *SYB* *SPRY* |  | *RAN* | *CYP* | *TBP1* | *EF2* | *EF1* | *MSF1* | *UBCE* | *EIF* | *MAPK* | *Actin* |
| NormFinder | *EF2* | *TBP1* | *EF1* | *SPRY* | *SYB* | *RAN* | *CYP* | *MSF1* | *EIF* | *MAPK* | *UBCE* | *Actin* |
| Bestkeeper | *SPRY* | *SYB* | *TBP1* | *RAN* | *EF2* | *EF1* | *CYP* | *MSF1* | *EIF* | *MAPK* | *UBCE* | *Actin* |
| **Comprehensive ranking** | *SPRY* | *SYB* | *EF2* | *TBP1* | *RAN* | *EF1* | *CYP* | *MSF1* | *EIF* | *UBCE* | *MAPK* | *Actin* |
| **Genorm** | *RAN MAPK* |  | *Actin* | *TBP1* | *SPRY* | *EF1* | *CYP* | *SYB* | *UBCE* | *MSF1* | *EF2* | *EIF* |
| **NormFinder** | *MAPK* | *RAN* | *Actin* | *TBP1* | *CYP* | *SPRY* | *EF1* | *SYB* | *EF2* | *UBCE* | *MSF1* | *EIF* |
| **Bestkeeper** | *Actin* | *RAN* | *MAPK* | *TBP1* | *SPRY* | *EF1* | *CYP* | *SYB* | *EF2* | *MSF1* | *EIF* | *UBCE* |
| **Comprehensive ranking** | *RAN* | *MAPK* | *Actin* | *TBP1* | *SPRY* | *CYP* | *EF1* | *SYB* | *UBCE* | *EF2* | *MSF1* | *EIF* |
| **Genorm** | *MAPK TBP1* |  | *EIF* | *SYB* | *EF2* | *Actin* | *CYP* | *UBCE* | *MSF1* | *RAN* | *SPRY* | *EF1* |
| **NormFinder** | *CYP* | *UBCE* | *Actin* | *EF2* | *MSF1* | *SYB* | *TBP1* | *MAPK* | *EIF* | *RAN* | *SPRY* | *EF1* |
| **Bestkeeper** | *SYB* | *TBP1* | *MAPK* | *EF2* | *EIF* | *Actin* | *CYP* | *UBCE* | *MSF1* | *SPRY* | *RAN* | *EF1* |
| **Comprehensive ranking** | *TBP1* | *Actin* | *EF2* | *SYB* | *CYP* | *MAPK* | *UBCE* | *EIF* | *MSF1* | *RAN* | *SPRY* | *EF1* |
| **Genorm** | *UBCE CYP* |  | *EIF* | *SYB* | *EF2* | *SPRY* | *MAPK* | *EF1* | *TBP1* | *MSF1* | *Actin* | *RAN* |
| **NormFinder** | *SPRY* | *EF2* | *SYB* | *UBCE* | *CYP* | *EIF* | *MAPK* | *MSF1* | *EF1* | *TBP1* | *Actin* | *RAN* |
| **Bestkeeper** | *SYB* | *SPRY* | *EF2* | *CYP* | *UBCE* | *EIF* | *MAPK* | *EF1* | *MSF1* | *TBP1* | *Actin* | *RAN* |
| **Comprehensive ranking** | *UBCE* | *SYB* | *CYP* | *EF2* | *SPRY* | *EIF* | *MAPK* | *EF1* | *MSF1* | *TBP1* | *Actin* | *RAN* |
| **Genorm** | *EIF Actin* |  | *SYB* | *EF2* | *MSF1* | *EF1* | *RAN* | *TBP1* | *SPRY* | *CYP* | *UBCE* | *MAPK* |
| **NormFinder** | *Actin* | *EIF* | *SYB* | *EF2* | *MSF1* | *EF1* | *TBP1* | *RAN* | *SPRY* | *CYP* | *UBCE* | *MAPK* |
| **Bestkeeper** | *EIF* | *MSF1* | *Actin* | *SYB* | *EF1* | *EF2* | *RAN* | *TBP1* | *SPRY* | *CYP* | *MAPK* | *UBCE* |
| **Comprehensive ranking** | *Actin* | *EIF* | *SYB* | *MSF1* | *EF2* | *EF1* | *TBP1* | *RAN* | *SPRY* | *CYP* | *UBCE* | *MAPK* |
| **Genorm** | *MSF1 SYB* |  | *MAPK* | *TBP1* | *SPRY* | *EIF* | *CYP* | *EF2* | *EF1* | *RAN* | *Actin* | *UBCE* |
| **NormFinder** | *MSF1* | *SYB* | *MAPK* | *EF2* | *TBP1* | *SPRY* | *CYP* | *EIF* | *RAN* | *EF1* | *Actin* | *UBCE* |
| **Bestkeeper** | *MSF1* | *CYP* | *SPRY* | *SYB* | *TBP1* | *MAPK* | *EF2* | *EIF* | *RAN* | *Actin* | *EF1* | *UBCE* |
| **Comprehensive ranking** | *MSF1* | *SYB* | *MAPK* | *TBP1* | *SPRY* | *EF2* | *CYP* | *EIF* | *RAN* | *EF1* | *Actin* | *UBCE* |

## Discussion

RT–qPCR is now commonly used in scientific research for the analysis of gene expression; thus, it is critical to select a suitable reference for improving the accuracy and reliability during qRT–PCR process. However, there was no consistent reference gene expression stability under different conditions suitable for all species. Therefore, a lot of studies have been conducted to select stable reference genes under certain conditions. In our study, a total of 12 selected reference genes were evaluated using a transcriptomic method and three algorithms. This is the first detailed study on the stability of internal control genes for RT–qPCR analysis in *P. portentosus* and revealed that transcriptome data could help to screen suitable genes under different conditions. The results will be beneficial to gene expression analysis in this fungus.

There are many genomic and transcriptomic datasets deposited in NCBI databases. These resources are an efficient way to mine and evaluate reference candidates [43–45,47]. Huge number of studies working on reference gene validation have been performed in plants (e.g., Arabidopsis, *Brassica napus*, and rice) and humans by selecting stably expressed genes from microarray and transcriptome datasets [8,11,39,52,53]. During this processing, 11 out of 25 traditional reference genes were maintained in this study. A total of 14 genes were excluded, including the commonly used internal control genes *18S* rRNA and *GAPDH*. *18S* rRNA and *GAPDH* are the least stable genes in plants, fungi and animals, especially in mushrooms, e.g., *O. sinensis, A. heimuer, F. filiformis, A. cornea, V. volvacea*, and *G. lucidum* [16,27,30,34,54]. The results of the evaluation of the retained genes preprocessed using transcriptomic data were better than those directly evaluated using RT–PCR and *geNorm, NormFinder* and *BestKeeper*. Based on genomic and transcriptomic datasets in *B. napus* L., a total of 12 genes could be used as appropriate reference genes and were better than traditionally used housekeeping genes [53]. In addition, some novel candidate genes could be mined using transcriptomic methods. In *Arabidopsis*, 8 newly discovered reference genes were mined identified using omics method [13]. In this study, the gene *SYB*, with the lowest CV value based on the transcriptome data during the developmental stages, was also selected. Validation using different algorithms showed that *SYB* was one of the most stable reference genes in line with the result of transcriptome. All the results revealed that pretreatment with multiple transcriptomic datasets could provide unimaginable information about gene expression variation across different stages and treatments and hence provide good reference gene candidates [55].

In this study, MSF1, *SYB, MAPK*, TBP1 and SPRY were the most stable reference genes when all the results generated from the different methods were combined. In *Volvariella volvacea, SPRY* and *MSF1* were the most stably expressed genes under different designed conditions [16], and *MAPK* was the best reference gene under acid treatment [31]. *TBP* (TATA-box-binding protein) was proved to be most stably expressed under different conditions and was sufficient for reliable results in humans [56]. In contrast, *EF1, Actin* and *UBCE* were unstably expressed in most tested samples of *P. portentosus. Actin* was the least stable and was deemed to be not suitable as an internal control for *V. volvacea, C. militaris*, and *L. edodes* gene expression studies under different developmental stages and in different media [14,16,17]. *ACT1* was the best internal control gene in *F. filiformis* [33,34]. The stability of *UBC* was not consistent with previous studies; *UBC* was one of the most stable genes in *Sacha inchi* seedlings [57], pearl millet [*Pennisetum glaucum* (L.) R. Br.] [58], Japanese flounder (*Paralichthys olivaceus*) [59], developmental and postharvested fruits of *Actinidia chinensis* [60], pepper (*Capsicum annuum* L.) [61] and *Eucommia ulmoides* Oliv [62]. All the results revealed that there were no consensus reference genes in all the species.

It was suggested that combination of two or more reference genes should be used for RT–qPCR analysis to generate more reliable results [63,64]. Pairwise variation calculated in *geNorm* could be used to determine how many reference genes were needed for an accurate analysis. When analyzing all experimental groups under different conditions, all had V scores much lower than 0.15, which indicated that there was no need to selection of combination different reference genes for normalization. However, the results must be in line with M values generated by *geNorm*. Because of no all the genes with M<0.2, two reference genes were adequate for RT-qPCR analysis. And MSF1 and *SYB* were selected as the best references for *P. portentosus* under different conditions.

## Conclusions

After evaluating the expression stability of 12 candidate reference genes using *geNorm, NormFinder* and *RefFinder* under different developmental stages and conditions, different genes were suitable for different conditions in *P. portentosus. MSF1, SYB, MAPK, TBP1* and SPRY were the top 5 stable reference genes in all sample treatments, while elongation factor 1-alpha (*EF1*), *Actin* and ubiquitin-conjugating enzyme (*UBCE*) were the most unstably expressed. Two reference genes were adequate for RT–qPCR studies to generate high accuracy results. The combination of *MSF1* and *SYB* might be the best selection. This is the first systematic study for selecting and validating reference genes in this fungus and provides guidelines for identifying genes and quantifying gene

## Supporting information

The amplified fragments of candidate reference genes shown by agarose gel electrophoresis

## Acknowledgments

This study was founded by the Science and Technology Innovation Action of Shanghai Science and Technology Commission (17391900400) and the National Natural Science Foundation of China (31800015).

## Data Availability Statement

All relevant data are within the manuscript and its Supporting information files.

