## Supplementary material for "Selection and validation of internal control genes for quantitative real-time RT–_q_PCR normalization of *Phlebopus portentosus* gene expression under different conditions": Table S1

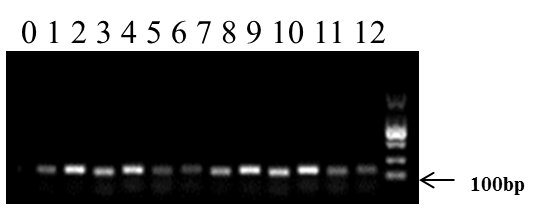

Figure S1. The Amplified fragments of candidate reference genes shown by agarose gel electrophoresis. 1-12 lanes: CK, *MSF1, SPRY, EF2, RAN, EIF, UBCE, EF1, MAPK, TBP1, SYB, Actin, CYP*
